# Improving replicability using interaction with laboratories: a multi-lab experimental assessment

**DOI:** 10.1101/2021.12.05.471264

**Authors:** Iman Jaljuli, Neri Kafkafi, Eliezer Giladi, Ilan Golani, Illana Gozes, Elissa J. Chesler, Molly A. Bogue, Yoav Benjamini

## Abstract

Experimentation with mouse and rat models has become a central strategy for discovering mammalian gene function, and for preclinical testing of pharmacological treatments, yet the utility of any findings critically depends on their replicability in other laboratories. In previous publications we proposed a statistical approach for estimating the inter-laboratory replicability of novel discoveries made in a single laboratory. We demonstrated that previous phenotyping results from multi-lab databases can be used to derive a Genotype-by-Lab (GxL) adjustment factor to greatly enhance the replicability of the single-lab findings, for similarly measured phenotypes, even before making the effort of replicating these finding in additional laboratories.

This demonstration, however, still raised several important questions that could only be answered by an additional large-scale prospective experiment: 1) Does GxL-adjustment work in single-lab experiments that were not intended to be standardized across laboratories, and with genotypes that were not included in the previous experiments? And 2) Can it be used to adjust the results of pharmacological experiments? We investigated these questions by attempting to replicate, across three laboratories, results from five single-lab studies in the Mouse Phenome Database (MPD), offering 212 comparisons, including 60 involving a pharmacological treatment: 18 mg/kg/day fluoxetine. In addition, we define and use a dimensionless GxL factor, by dividing the GxL variance by the standard deviation between animals within groups, as a more robust vehicle to transfer the adjustment from the multi-lab analysis to very different labs and genotypes.

For genotype comparisons, GxL-adjustment reduced the rate of non-replicable discoveries from 60% to 12%, for the price of reducing the power to make replicable discoveries from 87% to 66%. In absolute numbers, the adjustment prevented 23 non-replicable discoveries for the price of missing only three replicated ones. Tools and data needed for deployment of this method across other mouse experiments are publicly available in MPD. Our results further point at some phenotypes as more prone to produce non-replicable results, while others, known to be more difficult to measure, are as likely to produce replicable results (once adjusted) such as the physiological measure, body weight.

## 1. Introduction

The scientific community is concerned with issues of published results that fail to replicate in many fields including those of preclinical animal models, drug discovery, and discovering mammalian gene function (Freedman et al. 2015, Chalmers and Glasziou, 2009, Howells et al. 2014). Indeed, some of the first concerns regarding the complex interaction between genotype and the conducting laboratory were raised in the field of rodent behavioral phenotyping (Crabbe et al. 1999). While mouse and rat models may predict the human situation, such as the case of activity-dependent neuroprotective protein (ADNP) and the potential of its fragment as a drug (reviewed in Gozes, 2020), the utility of any findings critically depends on their replicability in other laboratories (Kafkafi et al., 2018; Voelkl et al., 2018). A similar concern arises regarding the interaction between the conducting laboratory and novel pharmacological treatments (e.g., Rossello et al., 2019) that are of vital importance for translational research, leading to novel drug developments.

In a previous publication (Kafkafi et al., 2017), we proposed a statistical approach for estimating the inter-laboratory replicability of novel discoveries at the level of the single laboratory study. This “random lab model” (Kafkafi and Benjamini, 2005) approach adjusts any phenotyping discovery in a single laboratory, by adding the Genotype × Laboratory (GxL) interaction “noise” σ^2^_G×L_ to the individual animal noise, thus generating a larger yardstick, against which genotype differences are tested and confidence intervals are reported. Consequently, this “GxL adjustment” raises the benchmark for discovering a significant genotype effect, trading some statistical power for ensuring replicability. We demonstrated (Kafkafi et al., 2017) that previous phenotyping results from multi-lab databases can be used to derive a GxL-adjustment term to ensure the replicability of single-lab results, for the same phenotypes and genotypes, even before making the effort of replicating the finding in additional laboratories.

This demonstration, however, still raises several important questions. Kafkafi et al. (2017) used the data from multi-lab, coordinated research programs to estimate the standard deviation of the GxL interaction, which was then used to adjust the results of each of these labs separately, as if the results from the other labs were unknown. While the success of this demonstration is encouraging, it does not cover the realistic setting where the adjusted laboratories are operating independently from the laboratories used for generating the GxL adjustment. Here, we therefore first investigate the question whether GxL adjustment reduces the proportion of non-replicable results, relative to the naïve analysis, and what loss of power does it involve.

A different yet important question is whether GxL estimation from standardized studies can be used to successfully identify replicable results in studies lacking special attention to standardization. Namely, will the adjustment based on the data from the International Mouse Phenotyping Consortium (IMPC, de Angelis et al., 2015; Muñoz-Fuentes et al., 2018), which typically uses relatively well-coordinated, standardized protocols, predict the replicability of results obtained in more common and realistic scenarios, such as those in the Mouse Phenome Database (MPD, Bogue et al., 2020). Unlike the IMPC, MPD archives previously conducted studies, which were not *apriori* meant to be part of a multi-lab project. Their methods, apparatus, endpoints and protocols of such experiments are thus not expected to be standardized.

Finally, our previous demonstration of GxL adjustment tested only genotype effects, using inbred strains and knockouts, but not pharmacological effects. It therefore remains to be tested whether the pre-estimated interaction of treatment with lab (TxL) or the interaction of the genotype and pharmacological treatment with the lab (GxTxL) can also be used to adjust single-lab treatment testing in a similar way.

In the present study we assessed the value of the GxL adjustment for experimental results previously submitted to the MPD involving genotype differences in several phenotypes. For that purpose, we conducted an experiment across three labs, without strong inter-laboratory standardization and coordination. In order to carry over the estimated interaction to a new laboratory, we scale the GxL interaction standard deviation (noted by σ_G×L_) by the pooled standard deviation between the animals within the groups (noted by σ), to define the unitless GxL factor γ = σ_G×L_/σ (termed environmental effect ratio by Higgins et al. 2021). The implication of this improvement is that rather than using the ‘raw’ σ_G_×_L_ for adjustment, we multiply it by the ratio of the standard deviation in the adjusted study to the one in the multi-lab study from which the interaction is estimated. The use of the scale-free dimensionless factor γ enables us to carry the information about the size of the interaction of a phenotype to other laboratories, other genotypes and variations in setups and conditions.

Using this 3-labs experiment we were able to test the replicability of the original results and show that using the GxL adjustments in the original studies could have greatly reduced the number of non-replicable discoveries.

## 2. Results

We conducted the phenotyping experiment (3-lab experiment) in the following three laboratories: the Center for Biometric Analysis core facility in The Jackson Laboratory, USA (JAX); the George S. Wise Faculty of Life Sciences, Tel Aviv University, Israel (TAUL); and in the Faculty of Medicine, Tel Aviv University, Israel (TAUM). We compare the effects of 6 mouse genotypes, and one pharmacological treatment (18 mg/kg fluoxetine) in several behavioral phenotypes, as well as in one straight-forward physiological phenotype (body weight). The original results in these comparisons were reported by 5 previous studies submitted to MPD. Most of these phenotypes were also measured in multiple labs in the IMPC, which could therefore serve as a second source for estimating the GxL factors, as in Kafkafi et al. (2017).

### 2.1 Replication in the 3-lab experiment

We first use our 3-lab experiment to evaluate whether the specific MPD studies’ results are replicable or not, using the Random Lab Model analysis. Therefore, Figures 1 and 2 (and Supplementary Figures S1–S5) display the 3-lab results and their summaries, together with the standard deviation of the within-group error, and the standard deviation of the GxL interaction. Consider for example Fig. 1 displaying the Body Weight (BW) phenotype results: each boxplot (top) displays the results for the corresponding genotype on the horizontal axis, in one lab identified by color, where three boxplots correspond to the three labs which are clustered together. When available for this genotype, a fourth boxplot displays the corresponding result from the original MPD study, in this case Crowley1 for females or Tordoff3 for males. The three panels display results for the females (left), the control males (center) and the fluoxetine-treated males (right). The bottom part of the figure displays the mean of each group, *σ_G×L_* and *σ.* In Fig. 1 the values in the bottom row are displayed in their original scale, but in other figures when a transformation was used for the analysis, the values are shown on the transformed scale. Note that there are some consistent differences between labs, expressed as vertical distances between lab lines, but these will not affect the replicability when comparing genotypes within the same lab (Kafkafi et al. 2018). The concern is rather the GxL interaction, which can be perceived by differences in slope across lab lines, while parallel lines represent no interaction. The black error bars represent the interaction standard deviation *σ_G×L_* and the within-group standard deviation *σ* (as in Kafkafi et al. 2005; Kafkafi et al. 2017). The ratio of the first to the second bars is the estimated GxL factor γ (see Statistical Methods).

**Figure 1.**
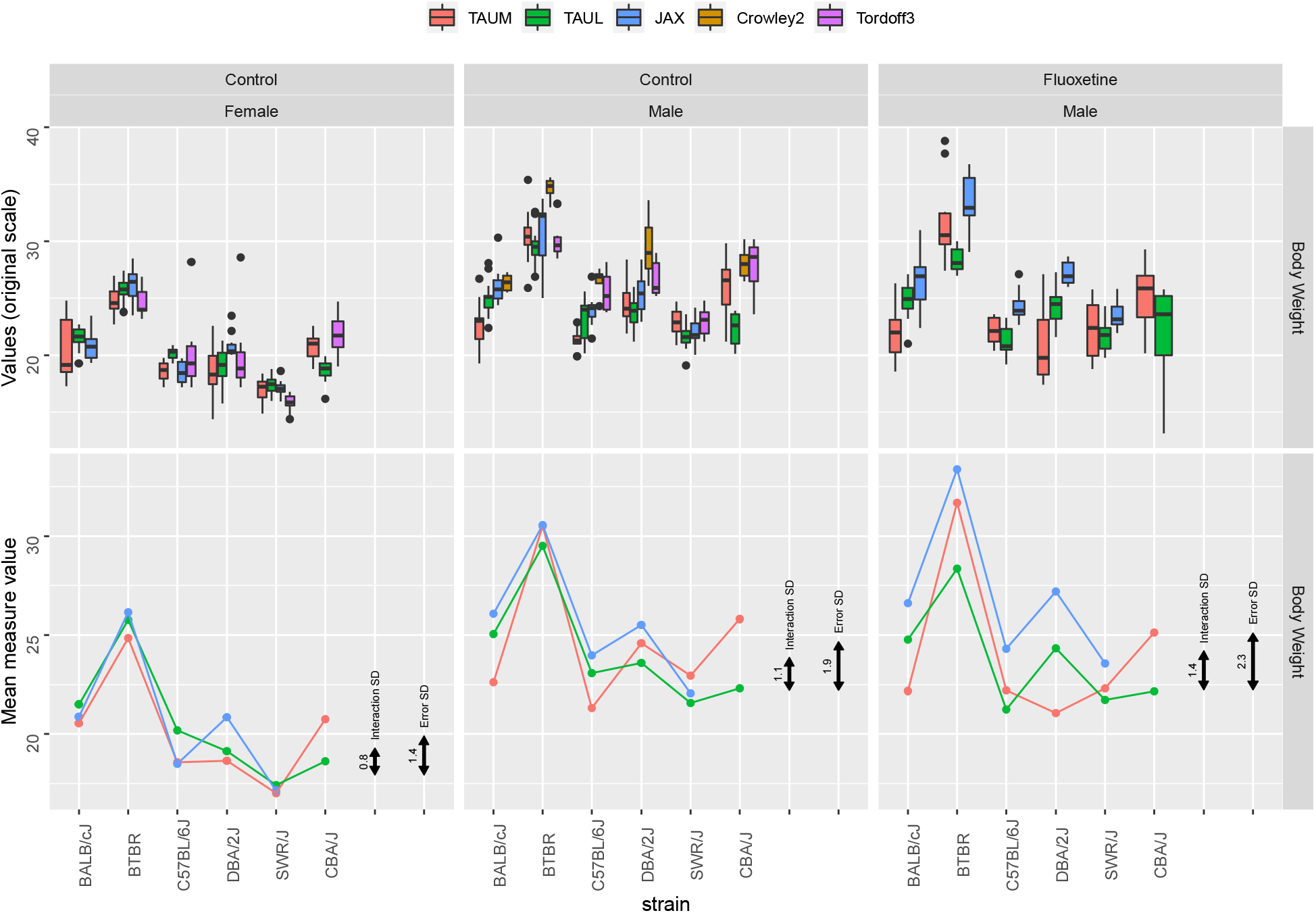
Body Weight (BW) in the 3 labs and in MPD studies Crowley1 and Tordoff3, using boxplots (top) and genotype means (bottom) in the three laboratories, in females (left), males (center) and fluoxetine-treated males (right). Black error bars represent the interaction SD and the within-group error SD.

**Figure 2:**
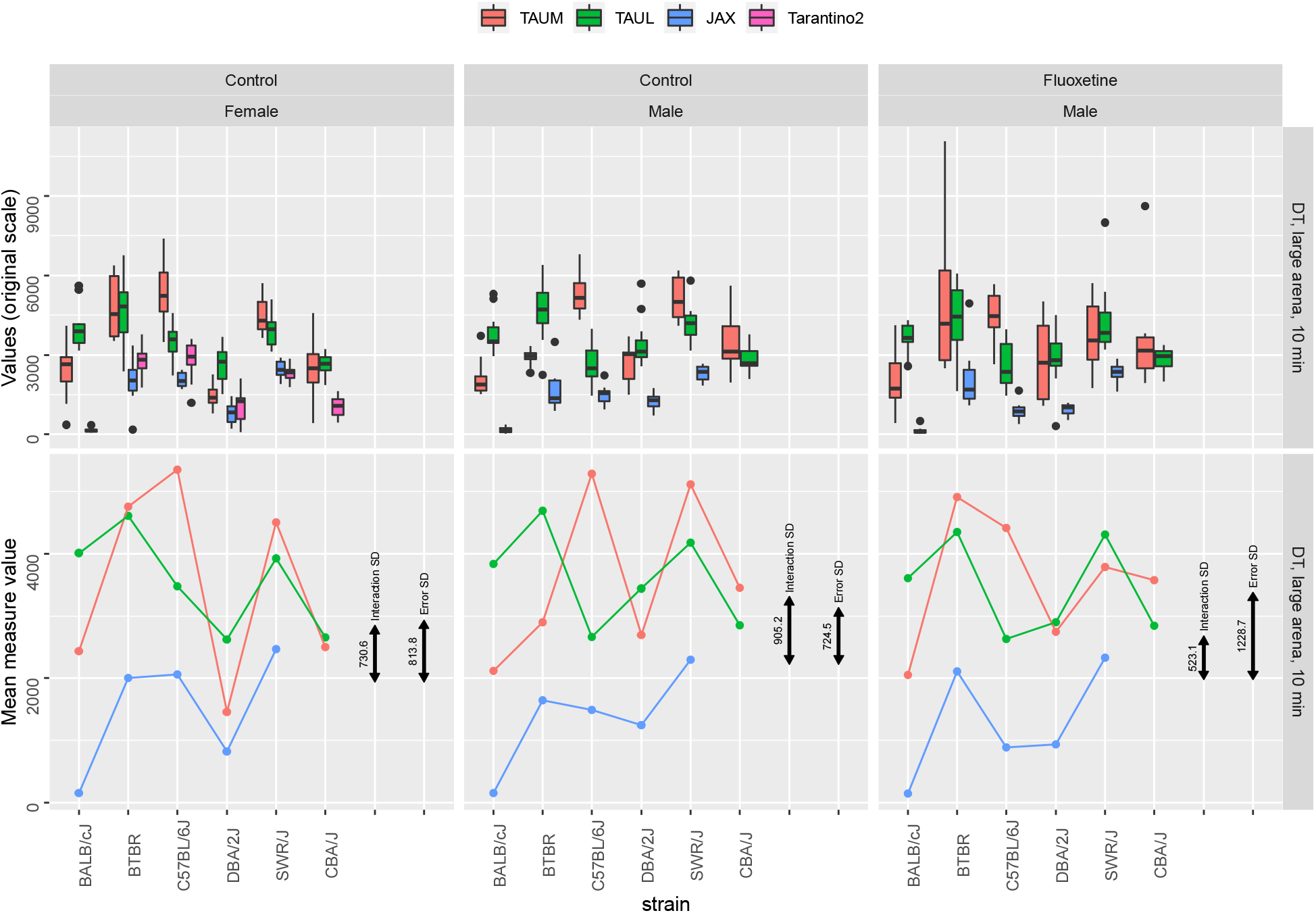
DT in the 3-lab, in a large arena for 10 minutes, and the MPD study Tarantino2 (for females). Graph organization is as in Figure 1, using boxplots (top) and genotype means (bottom) in the three laboratories: in females (left), males (center) and fluoxetine-treated males (right). Black error bars represent the interaction SD and the within-group error SD.

**Figure 3:**
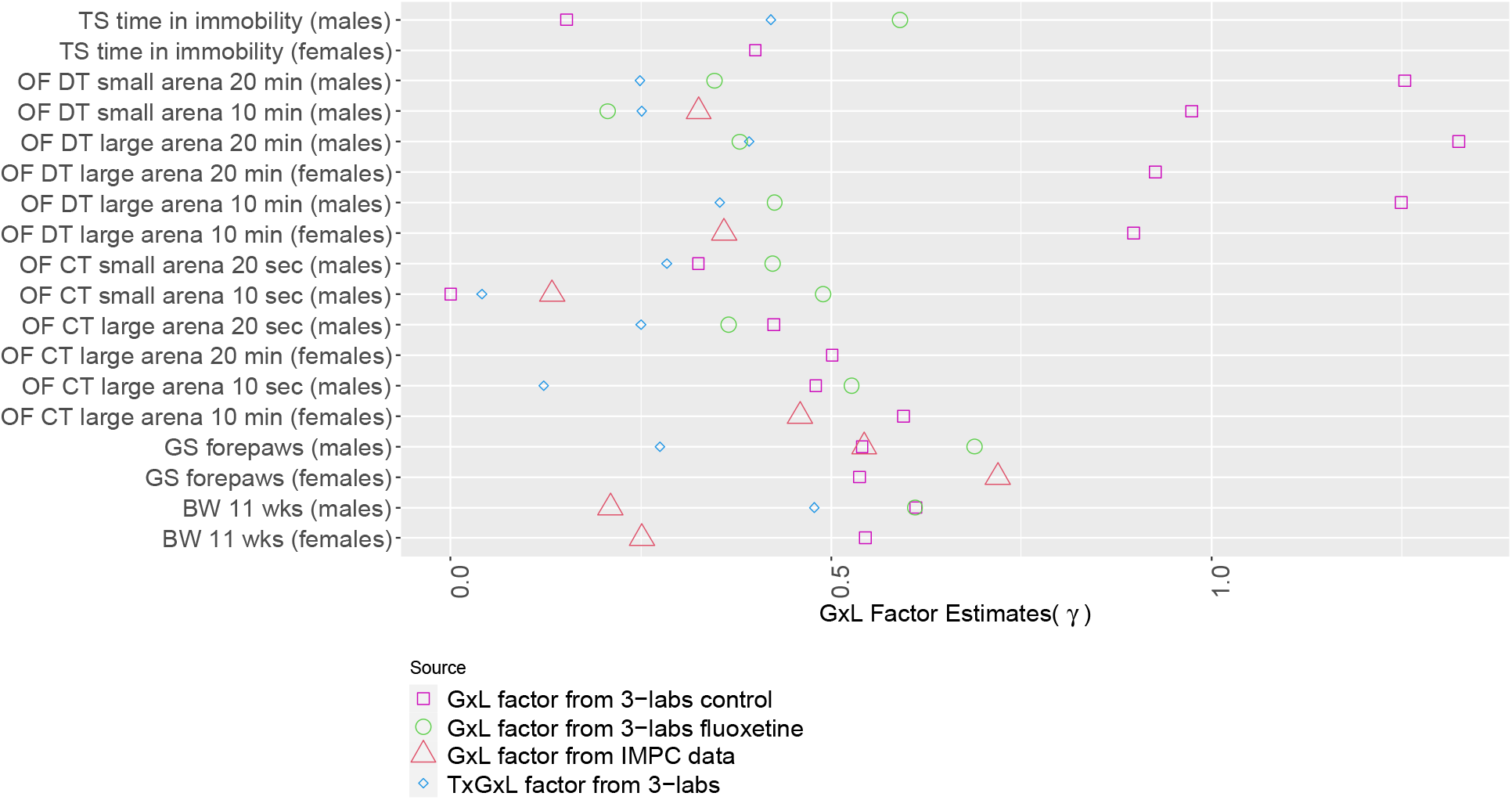
The values of the estimated interaction factor γ, for all endpoints as estimated from various sources: GxL factor from our 3-labs control data and from our fluoxetine treated data; GxL factor from IMPC data; TxGxL factor from our 3-labs data. CT and TS were logit transformed and GS was raised to the power of 1/3.

As expected from a reliably measured and well-defined physiological phenotype such as BW (Figure 1), the standard deviation of the error within the groups is small relative to the average weight, about 1.4/20 = 0.07 for females (left panel). Similarly, the interaction 0.8/20 = 0.04 is also relatively small. Yet the GxL factor, the ratio of these two standard deviations, is not negligible: *γ* = 0.8/1.4 = 0.57. In the fluoxetine-treated males (right panel), the GxL factor remains about the same *γ* = 0.61, although both the interaction standard deviation and the standard deviation of the error within groups increased, in fact by more than 50% (as evident from the bars).

In some behavioral phenotypes, notably the percent time spent immobile in the Tail Suspension (TS) test (Fig. S1), there were large lab absolute differences. This is hardly surprising, however, considering our use of different measurement technologies (force transducer method in JAX, as opposed to video tracking in TAUM and TAUL), as well as the choice of analysis parameters, such as the cutoff value for detecting immobility, which was also left for the specific lab to determine, as in typical single-lab studies. Despite this, the standard deviation of the interaction is still considerably smaller than that of the error within.

For the Distance Traveled (DT) endpoints, the estimated value of γ was near or larger than 1 (note especially Fig. 2 middle, where the interaction SD bar is actually larger than the Error SD bar). In contrast, all other phenotypes in this study had γ considerably smaller than 1 (see Figure 8 for summary). These high γ values do not appear to be the fault of any single genotype, lab, sex or treatment. Indeed, large γ occurred in all endpoints of DT, while they were still reasonable in the Center Time (CT) endpoints, which were measured in the same Open Field (OF) sessions (see discussion).

Note that in our experiment we could also measure related endpoints and variations in setups (such as 20 minutes session duration in OF, instead of 10 minutes) that were not required for adjusting an MPD experiment. We still estimated γ for them, teaching us about the robustness of the GXL interaction. These are all presented in Figure 8 and in Table S1.

### 2.2 GxL-Adjustment of phenotyping results

#### 2.2.1 GxL-adjustment of independent labs

The effectiveness of GxL-adjustment in Kafkafi et al., (2017) was not demonstrated for a phenotyping laboratory that is independent of the laboratories being used to estimate the GxL-adjustment. In this subsection we use the unitless GxL factor *γ*, estimated from the 3-lab experiment described in Section 2.1, the values of which are presented in Table S1 and displayed in Fig. 8. We examine the implications of the adjustment over all phenotypic differences between any two genotypes available (see “Statistical Methods” and Table 5.) Table 1 presents the results.

**Table1:** Results of naïve and GxL-adjusted genotypic differences for all phenotypes in the MPD experiments, using GxL factors estimated from the 3-lab.

| category | interpretation | No. cases | % cases (95% CI) |
| --- | --- | --- | --- |
| A | Replicable discoveries, approved by adjustment | 35 |  |
| B | Replicable discoveries, missed by adjustment | 11 |  |
| C | Replicable discoveries, missed by both | 7 |  |
| D | Non-replicable, insufficient adjustment | 12 |  |
| E | Non-replicable, prevented by adjustment | 47 |  |
| F | Non-replicable by both | 40 |  |
| (D+E)/(D+E+F) | Proportion of non-replicable discoveries | 59/99 | 59.6% (49-69) |
| (D)/(D+E+F) | Proportion of non-replicable discoveries, GxL-adjusted | 12/99 | 12.1% (6-20) |
| (A+B)/(A+B+C) | Power to discover replicable discoveries | 46/53 | 86.7% (77-94) |
| (A)/(A+B+C) | Power to discover replicable discoveries, GxL-adjusted | 35/53 | 66.0% (53-79) |

Out of the original studies’ significant findings, 60% (59/99) did not replicate in our 3-lab study. GxL-adjustment considerably decreased this proportion of non-replicable discoveries to 12% (12/99). The price paid in decreased power to detect replicable discoveries was a decrease from 87% (46/53) to 66% (35/53). In absolute terms, 47 non-replicable “discoveries” were prevented, while only 11 replicable discoveries were missed.

#### 2.2.2 GxL-adjustment using IMPC data

As in the previous section 2.2.1, in this section we use our 3-lab experiment for establishing the “ground truth”, where a statistically significant difference according to the Random Lab Model is considered as a “replicable discovery”. We also examine to what extent the number of non-replicable discoveries in an independent lab would have been reduced by adjusting it. However, unlike in 2.2.1 we use the GxL-interactions previously calculated from the IMPC multi-lab database (Kafkafi et al., 2017). The values of γ for the different endpoints are presented in Table S1 and displayed in Figure 8. Since IMPC database does not contain data of fluoxetine-treated mice, and does not include a TS test, the number of available genotypic differences in phenotypes is reduced to 92. Table 2 presents the results.

**Table 2:** Results of naïve and GxL-adjusted genotypic differences for all phenotypes in the MPD experiments, using GxL factors estimated from the IMPC standardized data.

| category | interpretation | No. cases | % Cases (95% CI) |
| --- | --- | --- | --- |
| A | Replicable discoveries, approved by adjustment | 29 |  |
| B | Replicable discoveries, missed by adjustment | 1 |  |
| C | Replicable discoveries missed by both | 3 |  |
| D | Non-replicable, insufficient adjustment | 14 |  |
| E | Non-replicable, prevented by adjustment | 16 |  |
| F | Non-replicable by both | 29 |  |
| (D+E)/(D+E+F) | Proportion of non-replicable discoveries | 30/59 | 50.8% (37- 63) |
| (D)/(D+E+F) | Proportion of non-replicable discoveries, GxL-adjusted | 14/59 | 23.7% (13-36) |
| (A+B)/(A+B+C) | Power to discover replicable discoveries | 30/33 | 90.9% (79-100) |
| (A)/(A+B+C) | Power to discover replicable discoveries , GxL-adjusted | 29/33 | 87.8% (76-97) |

**Table 3:** Results of naïve and GxLxT-adjusted genotypic differences for Wiltshire2 experiment, using GxLxT factors estimated from the 3-lab data. The rightmost two columns present results regarding differences in fluoxetine effect between genotypes, and the two columns left of them represent results regarding fluoxetine effect in single genotypes.

| category | interpretation | Fluoxetine effect in genotypes |  | Differences in fluoxetine effect between genotypes |  |
| --- | --- | --- | --- | --- | --- |
|  |  | No. cases | % cases (95% CI) | No. cases | % cases (95% CI) |
| A | Replicable discoveries, approved by adjustment | 1 |  | 0 |  |
| B | Replicable discoveries, missed by adjustment | 1 |  | 1 |  |
| C | Replicable discoveries missed by both | 2 |  | 8 |  |
| D | Non-replicable, insufficient adjustment | 1 |  | 0 |  |
| E | Non-replicable, prevented by adjustment | 2 |  | 1 |  |
| F | Non-replicable by both | 11 |  | 35 |  |
| (D+E)/(D+E+F) | Proportion of non-replicable discoveries | 3/14 | 21.4% (0-43) | 1/36 | 2.8% (0-8) |
| (D)/(D+E+F) | Proportion of non-replicable discoveries, GxL-adjusted | 1/14 | 7.1% (0-21) | 0/36 | 0.0% (0-10) |
| $(A+B)/(A+B+C)$ | Power to discover replicable discoveries | 2/4 | 50% (0-100) | 1/9 | 11.1% (0-33) |
| $(A)/(A+B+C)$ | Power to discover replicable discoveries, GxL-adjusted | 1/4 | 25% (0-75) | 0/9 | 0.0% (0-34) |

Using the IMPC-derived GxL-adjustment decreased the proportion of non-replicable discoveries in the original single-lab MPD studies from 51% to 24%, for the price in decreasing the power to detect replicable discoveries from 91% with no adjustment to 87%. In absolute terms, 16 non-replicable “discoveries” were prevented, while only 1 replicable discovery was missed. Comparing the adjustment offered by γ, as estimated from the standardized IMPC data as above, to the adjustment offered by γ estimated from our non-standardized 3-lab study, we notice that the latter tend to be larger (see Figure 8), as expected. In order to study the implication of this difference, we report the results of the analysis presented in 2.2.1 restricted to the set of differences reported here. The MPD based GxL-adjustment results in a proportion of non-replicable “discoveries” of 24%, while it is 10% when the adjustment is based on the 3-lab data. The power in the IMPC-based adjustment is 88%, in comparison to 61% using 3-lab-based adjustment (see Supplementary Table S2.)

### 2.3 Using GxL-adjustment for comparing drug-effects across genotypes

#### 2.3.1 Fluoxetine effect across six shared genotypes

Our 3-lab experiment shared six genotypes and three phenotypes with the Wiltshire2 study in the MPD, which estimated the effect of fluoxetine treatment. This study offered 15 treatment effects in genotypes and 45 pairwise comparisons of treatment effects between genotypes, where the “ground truth” could be derived from our 3-lab data, using a three-way random lab analysis with genotype-by-lab-by-treatment interaction. This interaction term is also relevant for adjusting single-lab comparisons of fluoxetine treatment between genotypes, so it could be estimated from the 3-lab experiment (see Statistical Methods.)

The number of results available for fluoxetine effect in genotypes is small, so the results of their adjustment should be viewed with much caution. The number of results for pairwise comparisons of fluoxetine effects between genotypes is larger, but most of them do not result in significant difference in the original study. Combining the two sets of results, the proportion of non-replicable discoveries was reduced from 8% (4/50) to 2% (1/50) by the adjustment, and the power to detect replicable discoveries was reduced from 23% (3/13) to 8% (1/13).

#### 2.3.2 Fluoxetine effect across 30 genotypes

The Wiltshire2 MPD study included 30 genotypes, between which 1305 pairwise comparisons of treatment effect can be conducted, with 45 of these included in our 3-lab experiment (already reported in Table 4). These 1305 comparisons can be used to assess the effect of the GxL-adjustment from 3-lab data for the three phenotypes. However, we do not have ground truth for these differences.

**Table 4:** Results of naïve testing and lower significance level of 0.005 genotypic differences for all phenotypes in the MPD experiments.

| category | interpretation | No. cases | % cases (95% CI) |
| --- | --- | --- | --- |
| A | Replicable discoveries, approved by adjustment | 15 |  |
| B | Replicable discoveries, missed by adjustment | 31 |  |
| C | Replicable discoveries missed by both | 7 |  |
| D | Non-replicable, insufficient adjustment | 3 |  |
| E | Non-replicable, prevented by adjustment | 56 |  |
| F | Non-replicable by both | 40 |  |
| $(D+E)/(D+E+F)$ | Proportion of non-replicable discoveries | 59/99 | 59.6% (50-69) |
| $(D)/(D+E+F)$ | Proportion of non-replicable discoveries, GxL-adjusted | 3/99 | 3.0% (0-7) |
| $(A+B)/(A+B+C)$ | Power to discover replicable discoveries | 46/53 | 86.8% (77 -94) |
| $(A)/(A+B+C)$ | Power to discover replicable discoveries, GxL-adjusted | 15/53 | 28% (17-41) |

**Table 5:** The results tables in the “Results” section use the same structure, categories and interpretation.

| “Ground truth”<br>by 3-Lab study | Single-lab<br>difference | Single-lab<br>GxL-adjusted | Category | Interpretation |
| --- | --- | --- | --- | --- |
| Replicable | significant | significant | A | Replicable discoveries, approved by adjustment |
| Replicable | significant | ns | B | Replicable discoveries, missed by adjustment |
| Replicable | ns | ns | C | Replicable discoveries missed by both |
| Non-replicable | significant | significant | D | Non-replicable, insufficient adjustment |
| Non-replicable | significant | ns | E | Non-replicable, prevented by adjustment |
| Non-replicable | ns | ns | F | Non-replicable by both |
|  |  |  | (D+E)/(D+E+F) | Proportion of non-replicable discoveries |
|  |  |  | (D)/(D+E+F) | Proportion of non-replicable discoveries, GxL-adjusted |
|  |  |  | (A+B)/(A+B+C) | Power to discover replicable discoveries |
|  |  |  | (A)/(A+B+C) | Power to discover replicable discoveries, GxL-adjusted |

For 276 comparisons, the regular t-based linear contrasts for fluoxetine treatment effects found significant difference in Wiltshire2 data. Two hundred four of them (73%, 95%CI 68.5-79.0) would become non-significant once the GxL-adjustment is used. The adjustment thus weeds out a large proportion of apparently significant differences, but we cannot tell if it is justifiable or not.

### 2.4 Lowering the significance threshold in the original analysis

One potential response to our previous results is that by incorporating the GxL-adjustment we have merely lowered the alpha-level of the test being used, and one can instead use a single lower level across all comparisons. We have tested the across-science recommendation by Benjamin et al (2018) to use the 0.005 level for statistical significance in the original studies. Table 4 shows the rejection rates when adapting the uniformly more conservative threshold.

Using the lower α = 0.005 indeed offers a more conservative test, where the proportion of the non-replicable discoveries is reduced to 3%, compared to the 12% using GXL-adjustment. However, this comes with a very large price in power, which is reduced to 28%, comparing to 66% using the GxL-adjustment.

## 3. Discussion

### 3.1 The contribution of GxL-adjustment to replicability

What performance can be expected from GxL-adjustment in realistic situations of single-lab studies, which are often not standardized with other labs? The most direct answer to this question is given in Section 2.2.1. Using database-derived interaction estimates for body weight (BW), distance traveled (DT) and center time (CT) in the open field test, grip strength, and tail suspension, the GXL-adjustment reduced the number of false discoveries by half, from 60% to 12%. This came with smaller loss of power, from 87% to 66%. In absolute terms, 47 non-replicable “discoveries” were prevented, while only 11 replicable discoveries were missed. It is important to emphasize that the original studies used only a few of the genotypes from which GxL-interactions were estimated. The testing parameters and conditions of testing were also somewhat different in both.

It might be argued that our success is merely due to our lowered thresholds for significance, and lowering the significance threshold to .005, as suggested in Benjamin et al., (2018) and its many signed supporters, will have similar results. However, as shown in Section 2.4, when implementing this suggestion for the above set of MPD original results, the result was so conservative that the power was 27%, instead of over 60% with GxL-adjustment. The GxL adjustment takes into consideration the different levels of adjustment needed for the different endpoints when facing the multi-lab replicability challenge, while using a single lower statistical discovery threshold across all phenotypes ignores the nature and robustness of each individual phenotype. Clearly there is an advantage to offering differing yardsticks to the different endpoints.

It should be emphasized that in order to isolate the effect of GxL adjustment on replicability we have treated each original result as though it were individually generated, ignoring any selection effect of the statistically significant ones among the many ones tested. Analyzing the structure of each original experiment and controlling the false discovery rate in the original studies would have further reduced the number of non-replicated results, below 12% but this is beyond the goals of the current study.

### 3.2 GxL adjustment by standardized multi-lab data

By making use of the IMPC data to estimate the GxL factor, we manage to stage the most realistic setting for checking the GxL adjustment in regular experimental work. The three tasks of i) generating the GxL-adjustment; ii) establishing the multi-lab “ground truth”; and iii) estimating the performances of the adjustment in single-lab studies, are conducted each in an independent set of laboratories: by the IMPC multi-lab data, by the 3-lab experiment, and by the MPD data, respectively. Unfortunately, using IMPC data for this purpose has an inherent limitation: The standardized way by which this multi-lab study is conducted, might not reflect in full the variability between typical labs that may differ in the protocol being used, its execution, and local conditions. This regular variation should be captured by the GxL factor, and not the lower variation yielding the smaller GxL factor. Thus, it is not clear whether the proportion of non-replicable discovery rate of 24% we get using the IMPC based adjustment, which is higher than the 10% we get using the 3-lab data for adjustment, reflects the more realistic setup or the less realistic source of data. Nevertheless, even if the actual implementation of our approach bounds the proportion of non-replicable results at somewhere between 24% and 10%, it far more comforting a number than the 61% offered by doing nothing.

These results further indicate that extending the current effort in the MPD to utilize varied experimental results, created under no special standardization, for estimating the GxL-adjustment, should yield better results than merely relying on a single large initiative such as IMPC. Because the breadth of phenotyping in the IMPC was necessarily limited, while the breadth of archival experiments is potentially unlimited, the use of a database of research results as in MPD is a promising approach for evaluating replicability across a wide range of experiments. MPD houses thousands of well-curated physiological and behavioral phenotypic measures, and each is stored with detailed protocol information that will allow users to choose a collection of data sets that has relevant procedural, environmental and genotypic characteristics for estimation of GXL. By coupling the GXL replicability estimator to this database, we have enabled users a facile means of evaluating the replicability of their findings and contributing data to future users wishing to do the same. The utility of the approach grows as the breadth and depth of the data resource is expanded. Global analyses of replicability within the MPD can inform the refinement of phenotyping paradigms in many areas of research.

### 3.3 GxL-adjustment of drug-treatment discoveries

A practical limitation of our effort to experimentally verify the utility of the GxL-adjustment for drug treatment experiments was the small number of results that were available for testing our approach. Indeed, the changes were small: there were only four strain differences in which the original and the adjusted analysis differed. However, relying on the analysis in Section 2.2.2, which offered hundreds of potential differences, by including many more genotypes, using the adjustment weeds out a large number of original discoveries. Of course, we have no way to verify the ground truth for genotype differences that were not tested in the 3-lab experiment, so performance based on the number non-replicable discoveries prevented and decreases in statistical power cannot be estimated, but the impact should be large. Reassuringly, the GxL factor estimated for fluoxetine for many phenotypes was larger by merely 5% than the value for control mice, and the 3-way interaction was close to that value. These two results suggest that it may mostly be a property of the phenotype used. Future work should establish whether the interaction of Lab-by-Drug-by-Genotype does not depend critically on the drug administered.

Our results should not be used to draw conclusions about the clinical efficacy of fluoxetine, since the traditional TS test does not necessarily predict anti-depressant efficacy in humans (Hyman 2012) and is therefore only used here to test the replicability of previously-published results in mice, which were available for us in MPD.

### 3.4 Identifying and restructuring problematic phenotypes

It follows from our work that for an endpoint to be useful, its design should take into consideration the size of its GxL factor *γ.* This factor compares the interaction variability to the animals’ variability, a point of view that may be different from current thinking. Body weight has low variability among animals, i.e., high precision, but its interaction term measured here was high. At the same time, tail suspension is notoriously known to be of high variability, but surprisingly the interaction is small (see also Fig 5 central column), and thus the ratio turned out close to that of body weight.

More surprisingly, in the common OF test, CT had a consistently smaller factor than DT. Indeed, DT has had the largest number of comparisons changed from replicable to non-replicable due to the adjustment. Interestingly, DT as well as the CT proved highly replicable in a previous OF test by some of us, in 8 inbred strains, some of them used in the current study, across three laboratories, all different than the laboratories in the current study (Kafkafi et al., 2005). Both the interaction and the Error SD were considerably smaller relative to the measured size, with *γ* ≈ 0.7 (see Figure 1 and error bars in Kafkafi et al., 2005). However, this previous study was conducted in circular, much larger arenas (≈ 250 cm diameter vs ≈ 40 cm width in the current study), while using standardized video tracking systems, and employing standardized SEE analysis for robust path smoothing and segmentation (Drai et al., 2001; Kafkafi et al., 2003). The DT, being a measure of change in location across time, is probably more sensitive than CT, a measure of location, to lab-specific tracking noise that depends on the tracking technology and parameters used in each laboratory (Hen et al. 2004).

Lipkind et al. (2004) demonstrated the use of multi-lab results to explicitly improve the design of the DT and CT endpoints with robust methods to achieve better replicability across laboratories. Thus, while Wurbel et al. (2021) recently argued against using behavioral tests, our stand is that behavioral testing should not be dropped but rather improved, not by specifying to finer resolution how the test should be conducted, but by directed design of test hardware and software for higher replicability.

### 3.5 Conclusion

In the present study across three laboratories, we explored 152 comparisons between mouse genotypes. Of course, not all of them are expected to reflect real differences, but 53 of them did turn out to be replicable, in the sense that they were significant in random lab model analysis across three independent labs. This indicates that, despite the criticism expressed at preclinical research using animal models, there are replicable signals worthy of exploration. Moreover, 46 of these 53 (87% power) were already discovered by the original single-lab studies. Unfortunately for the field, the criticism is correct in expressing alarm over the rate of non-replicable discoveries that comes with such a high power: along with the 46 replicable discoveries, the single lab studies also “discovered” 59 non-replicable ones.

Two solutions are generally offered for this unacceptable situation. The first one is increasing the sample size, as argued by Szucs and Ioannidis (2017). However, our work demonstrates the limitation of this recommendation for preclinical studies: it further increases the already sufficient power in the single lab, magnifying the impact of local peculiarities by making more of them statistically significant, while the within-group variance remains the same. These peculiarities will disappear relative to the much larger interaction variability that does not decrease with increasing sample size, making the additional findings non-replicable.

A second offered solution is conducting preclinical experiments across several labs, accepting only discoveries that pass the random lab analysis, as simulated by Voelkl et al. (2018) using data in the literature, and demonstrated here in our 3-lab analysis. A similar conclusion was reached by Schooler (2014) in the field of experimental psychology, where he advocated independent replications across laboratories before publication, and made the commitment to conduct his future work this way. However, while this multi-lab solution does work in principle, it also raises major practical difficulties for the explorative investigator: Convincing additional laboratories to participate before any findings have been published seems to be one such obstacle, and the larger budgets required are a second obstacle. In that same work, Schooler (2014) acknowledges that “it is clearly not feasible for all researchers to follow this approach in their routine work”, and indeed, even in his own work this remained an ideal not too often reached. Not the least important, a third obstacle particular to preclinical animal studies is that more animals need be sacrificed before there is an indication of a replicable and important result.

This problem therefore led to two similar and more practical approaches in the field of preclinical animal models. Richter et al (2001) suggest heterogenization of the setup in the single lab experiment in order to capture the variability of a multi lab study. While helpful, not all the multi-lab variability was indeed captured in that study (and see also the reanalysis in Kafkafi et al (2017)). Simulations of multi-lab data in Voelkl et al. (2018) give a more promising point of view and report better success although it is not quite clear what aspects of the experiment should be heterogenized in a single-lab scenario.

The second approach, the GxL adjustment method, has been shown here to be a good surrogate for such multi-lab experiments, solving almost entirely the problem of too many non-replicable discoveries. Future cooperation of scientists in the area to enrich the publicly available databases such as those reported in MPD, where GxL factors for new endpoints can be estimated, as well as investing efforts to design more replicability enhancing measurement tools in the sense of having lower GxL factors will enable preclinical research to benefit from experience and results from prior animal studies.

## 4. Materials and Methods

### 4.1 Databases and replicated studies

Two phenotypic databases are employed in this study: the Mouse Phenome Database (MPD, Bogue et al. 2020) and the International Mouse Phenotyping Consortium (IMPC, Muñoz-Fuentes et al., 2018). MPD includes previous single-lab mouse studies submitted by data contributors, and here we attempt to replicate across three laboratories some of the results in these experiments. This tool is now implemented in MPD: https://phenome.jax.org/replicability. The IMPC data was used to estimate interaction terms across several IMPC centers, as in our previous study (Kafkafi et al. 2017).

Results from five independent studies in the MPD include four tests that were chosen to be replicated (MPD study code names as they appear on the MPD website): Wiltshire2 (Benton et al., 2012): Open-Field (OF), Tail-Suspension (TS); Tarantino2 (Schoenrock et al., 2016): OF; Crabbe4 (Crabbe et al., 2003): Grip Strength (GS); Tordoff3 (Tordoff et al., 2007): Body Weight (BW); Crowley1 (Crowley et al., 2010): BW.

Several considerations led us to select these studies: while searching the MPD for studies comparing many genotypes on many phenotypes, we nevertheless had to limit the number of animals being tested in our 3-lab experiment, by testing each mouse for several phenotypic endpoints. We therefore looked for studies in the MPD which: (i) shared the same genotypes and sexes; (ii) shared the same phenotypes; (iii) the phenotypes were also limited to those for which data from IMPC was available for the interaction terms; (iv) maximize the number of statistically significant findings in the MPD studies, since these are the only ones that might potentially be refuted by the GxL-adjustment. Still, whenever selecting several phenotypes and genotypes, many differences were not statistically significant in the original studies.

### 4.2 Laboratories

The three labs replicating the MPD studies were: The Center for Biometric Analysis (CBA) core facility The Jackson Laboratory, USA (JAX) under Bogue’s supervision; The Laboratory in The George S. Wise Faculty of Life Sciences, Tel Aviv University, Israel (TAUL), under Gozes’ supervision; The laboratory in the Faculty of Medicine, Tel Aviv University, Israel (TAUM) under Golani’s supervision. At JAX mice were housed in the CBA animal room and testing was conducted in CBA procedure rooms; In TAUL, mice were housed in the Faculty of Life Animal House, and test were conducted by NK in the behavioral room in this facility. In TAUM, the animals were housed in David Glasberg Tower for Medical Research and tests were conducted by Eliezer Giladi (EG) on the 6^th^ floor of the Tower at the Myers Neuro-Behavioral Core Facility.

Note that the two labs in Tel Aviv University were in separate faculties and buildings, had separate experimental animals, facilities and technicians, and worked in independent time schedules. We took special care not to coordinate these two laboratories, as if each of them conducted the experiment in an independent study. Veterinary and Animal Care inspections in these two laboratories are both conducted by the TAU Center for Veterinary Care, but no active interference was required. All animal procedures in TAU were approved by TAU Institutional Animal Care and Use Committee and the Israeli Ministry of Health. At JAX, testing was performed in accordance with protocols approved by The Jackson Laboratory Institutional Animal Care and Use Committee.

### 4.3 Animals and Drugs

All three labs used the inbred strains: BALB/cJ, BTBR T^+^ Itpr3^tf^/J (BTBR), C57BL/6J, DBA/2J, SWR/J. The strain CBA/J was also used at TAUM and TAUL, but not at JAX. Breeders were transported from The Jackson Laboratory to each TAUM and TAUL and were distributed to cages of one male and 2-3 females. 1 to 4 litters from each breeder cage were then separated at ages of approximately 50 to 60 days old, to cages of 2-5 mice of the same strain and sex. For the JAX experiment, mice were shipped from The Jackson Laboratory’s production facility directly to JAX CBA and mice were identified by ear punch. In TAUL, mice were transferred to smaller cages in a different room than the breeders and were identified by tail marks. Half of the male mice were administered 18 mg/kg/day fluoxetine in drinking water (see below). Males were treated with fluoxetine, while females were not, because there were no such MPD experiments in females that we could attempt to replicate. Mice were tested at similar ages (see below).

Fluoxetine HCl was purchased from Medisca Inc., Lot 172601 (Plattsburgh NY USA) for the JAX experiment. In TAU, commercial fluoxetine HCl in 20 mg capsules was purchased from Ely Lilly, Israel. As in Wiltshire2 (Benton et al., 2012), the average weight measurements for each strain, together with previously determined daily water intake for each strain, were used to determine the amount of fluoxetine required to provide a daily oral dose of 0 or 18 mg/kg/day per mouse in drinking water. Male mice were treated daily with fluoxetine or water throughout the experiment. Fluoxetine treatment started at 5-6 weeks of age, in order to ensure three weeks of treatment before testing. In TAUM, the content of the water bottles changed every three days, while in TAUL and JAX they were changed every week, due to the use of larger bottles.

### 4.4 Tests, phenotypes and testing parameters

This study approximately followed the IMPC behavioral tests and protocols of Open filed (OF), Grip Strength (GS) and body weight (BW), from the behavioral pipeline of the IMPC IMPReSS EUMODIC pipeline 2 https://www.mousephenotype.org/impress/PipelineInfo?id=2 (Ayadi et al., 2012). In addition, we replicated the Tail Suspension Test in Wiltshire2 (Benton et al., 2012), which is not included in the IMPC pipelines (see statistical methods).

The interaction terms of genotypes with the laboratory were previously estimated across multiple laboratories by Kafkafi et al. (2017) and were also measured in experiments submitted to MPD (see “Databases” above). Experiments and genotypes were chosen to maximize the number of tests and phenotypes with previous interaction terms from multiple labs, as explained in the considerations below. The OF test included the phenotypes (Table 1) of Distance Traveled (DT) and the percentage of time spent in the center (CT). The TS test measured the percentage of time spent immobile, and the GS test measured forepaws peak grip strength, using the average of three consecutive measures.

Due to the differences between the IMPC pipeline and the different MPD studies, as well as local constraints in the three labs, it was not feasible to precisely standardize the identity of tests, their order and the ages in which they were conducted. Indeed, such precise standardization does not represent the realistic situation of the field and is unsuitable to the objective of this study.

However, age differences at the time of each test were at most 5 weeks, and all mice were post-pubertal and relatively young adult ages, i.e., not middle aged (12 month) or aged (≈ 18 + months). Table S1 summarizes the timelines in the databases, replicated MPD studies and the three labs. For similar reasons, the parameters and conditions of each test were not precisely standardized. These differences are detailed below, and for the OF test are also summarized in Table S2.

In the OF test, the phenotypes of distance traveled and percent of time spent in the center, in a small arena, for 10 and 20 minutes were recorded. In TAUL and TAUM these were also measured in a large arena (Table S2).

#### Open Field (OF) methods

Mice were allowed to acclimate to the testing room for a minimum of 60 min. Arena parameters were slightly different in the IMPC database, MPD studies and the three replicated labs, and are summarized in Table S2. The apparatus was a square chamber, either 27 x 27 cm (“small”) for males (as in Wiltshire2) or 40 x 40 cm to 50 x 50 cm (“large”) for females (as in Tarantino2 and the IMPC protocol). In TAUL and TAUM, the males were also tested in a large arena, a week after all the other tests were concluded (Table S2) in order to facilitate comparisons with the IMPC results.

Center and periphery definitions were also different (Table S2). The Session duration was 20 min (as in the IMPC protocol), but all analysis was done for the first 10 minutes (as in Wiltshire2) as well. In each test, the total distance traveled (DT) and the percentage of session time spent in the center of the arena (CT) were measured. In addition, TAUL and TAUM also tested the control and fluoxetine males in a second OF session in the “large” arenas (as in the IMPC database) about a week after completing all other tests. Between subjects, the arena was cleaned with 70% ethanol. In TAUL, mice were tested four at a time in four square Plexiglas arenas. To begin each test, the mouse was placed in the center of the arena. The apparatus was a square chamber either 27 x 27 cm or 50 × 50 cm. Tracking and Analysis was conducted using a Noldus EthoVision video tracking system. In TAUM, mice were tested 4 at a time in 4 square Plexiglas cages. To begin each test, the mouse was placed in the center of the arena. The apparatus was a square chamber either 27 x 27 cm or 50 × 50 cm. Video Tracking and Analysis was done with Noldus EthoVision video tracking system. In JAX, mice were acclimated to the testing room for a minimum of 60 min. The apparatus (Omnitech Electronics, Inc., Columbus OH, USA) was a square chamber (40 × 40 cm). To begin each test, the mouse was placed in the center of the arena. Data were recorded via sensitive infrared photobeams and collected in 5-min bins.

#### Grip Strength (GS) methods

Three trials were carried out in succession measuring forelimb-strength only and averaged, as in the replicated MPD experiment Crabbe4 (Crabbe et al., 2003) and the IMPC protocol. The mouse was held by the tail, lowered over the grid, keeping the torso horizontal and allowing only its forepaws to attach to the grid before any measurements were taken. The mouse was pulled gently back by its tail, ensuring the grip the on the top portion of the grid, with the torso remaining horizontal. The testing area was cleaned with 70% ethanol between subjects. In TAUL, TSE Systems Grip Strength Meter for mice was used with a mesh grid. In TAUM a commercially available Ugo Basile Grip-Strength Meter was used with a wire grid, coupled with a strain gauge measuring peak force in kg. In JAX, mice were acclimated for 60 min prior to testing. A commercially available grip strength meter (Bioseb, Pinellas Park FL, USA) was used.

#### Tail Suspension (TS) methods

Mice were allowed to acclimate to the testing room for a minimum of 60 min prior to testing. They were suspended by their tails with adhesive tape to the top of Plexiglas cages. The percentage of time spent immobile was measured in 6 and in 7 minutes. Between subjects, the testing area was cleaned with 70% ethanol. We used the 7 min data, as did Wiltshire2 (Benton et al., 2012) who reported dropping the first 1 minute because all mice remained mobile throughout. This led to not dropping the last minute in our results and should hardly affect differences. In TAUL, Tracking and Analysis was measured with Noldus EthoVision video tracking system. Polyethylene cylinders about 24 mm tall and 10 mm in diameter on the base of the tail were used to minimize the ability of mice to climb on their tails. Several mice that did manage to climb on their tails were discarded from analysis, as in the original Wiltshire2 experiment (Benton et al., 2012). In TAUM, Tracking and Analysis was measured with Noldus EthoVision video tracking system. In JAX, standard Med-Associates (St. Albans VT, USA) Tail Suspension Test chambers were used. Mice (10-12 weeks) were suspended by their tails with adhesive tape (VWR or Fisher, 25 mm wide) to a flat metal bar connected to a force transducer that measures transduced movement. The tape was extended the length of the mouse’s tail from 2 mm from base through 2 mm from tip, minimizing the ability of the mouse to climb its tail. A computer interfaced to the force transducer recorded the data.

#### Number of animals

We designed the experiment for at least n=10 mice per group. In some groups the number of animals was larger, due to litter sizes. In 10% (52/505) of the groups there were 9 animals. However, in tail suspension testing in JAX, some mice climbed their tails and no measurements were available, resulting in 4 measurements per group in four groups of females; 6 measurements per group and 8 measurements per group in 4 and 3 groups of males respectively. In 6% of the groups there were 15 or more animals. In the MPD experiments the number of mice per group ranged from 5 to 20, with median of 10.5 and interquartile range of 5. In the IMPC data the number of mice per group ranged from 3 to 403, with a median of 8 and interquartile range of 3. We obviously had no control of the number of mice in MPD and IMPC.

### 4.5 Statistical Methods

#### 4.5.1 GxL-adjustment of genotype effect in a single-lab study

The conventional t-test for testing phenotypic difference between genotype x and genotype y, is:

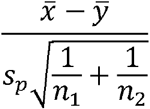

where n_1_ is group size for genotype x and n_2_ for genotype y, and *s_p_* is the pooled standard deviation within the groups. The number of degrees of freedom is *df* = *n*_1_ + *n*_2_ − 2. The underlying assumptions are that both genotype groups are independent, that both have equal within group variances, and that the distribution of the phenotype is approximately Gaussian. Appropriate transformations of the original measurements such as log, logit and cube root were used to make these assumptions more appropriate. The test can be modified if the variances are grossly unequal in the two groups.

##### The Random Lab Model for replicability

When a phenotype is compared between G genotypes in L laboratories, Kafkafi et al (2017) introduced the existence of Genotype by Lab interaction (GxL) as a random component the variance of which is 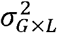. The implication is that:

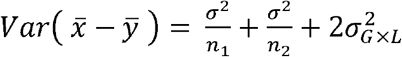

This alters the t-test as follows:

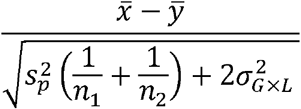

##### GxL-adjustment from a database

We define the GxL factor *γ* estimated from the results of a multi-lab study or database, as the square root of the ratio of the interaction variance to the pooled within-group error in the variance:

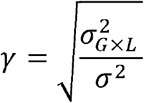

Note that *γ*^2^ is the environmental effect ration (EER) of Higgins et al. (2021)

When using GxL-adjustment at the single lab, the t-test, with its estimated standard deviation *s_p_*, becomes

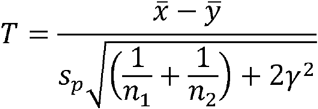

The underlying assumption is that this single-lab study comes from the same population as the multi-lab study, but the local measure may have a different multiplicative scale which is reflected by the ratio of the within group standard deviations. Hence 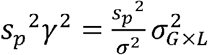 is an estimator of the interaction in the new study. Multiplying the database estimated interaction term by the ratio of the standard deviation in the adjusted single lab to the pooled variance in the multi-lab database translates the interaction term in one set of labs into a more relevant one in the single lab.

In a pre-clinical experiment involving laboratory mice and rats, a typical batch size is of 10–20 animals, hence, 1/*n_i_* is typically smaller than 0.1. This gives us an interpretation of the size of *γ*^2^: *γ*^2^ ≈ 0.1, namely *γ* ≈ 0.3, is of the same order as 1/*n*, and *γ*^2^ > 1 would have a very large effect. Note that the group sizes *n*_1_, *n*_2_ do not have any effect on the GxL factor *γ.* That is, if the interaction variance is large, increasing the number of animals can hardly improve replicability. The distribution of T is approximated by Student’s *t* with *v* degrees of freedom using the Satterthwaite formula: Let *n_L_* and *n_s_* denote the number of labs, and genotypes used for estimating *γ*^2^

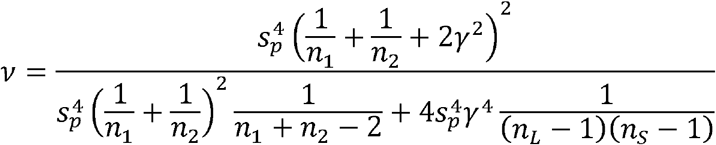

#### 4.5.2 Using GxL-adjustment for treatment effect

The database which we use for estimating the adjustment size *γ*^2^ only offers it for tests where animals were not treated, although tests for treatment effect are an important part of animal testing. Note that the adjusted test statistic T is still valid in cases where both groups are treated with similar treatment. Therefore, we also use it to test the replicability of strain effect when both groups are treated with fluoxetine, even though the adjustment component was estimated via untreated animals.

Multi-lab experiments that also included treatment groups are analyzed using a random-lab 3-way analysis, where treatment effect is added as a fixed effect, in addition to the factors we originally have in the random-lab 2-way analysis. We also include interactions involving treatment of all orders, namely, treatment-by-genotype, treatment-by-lab, genotype-by-lab and treatment-by-genotype-by-lab where the proportional SD of the latter interaction to error SD is denoted by *γ_T×G×L_*.

In their paper (Benton et al., 2012) Wiltshire2 test the effect of fluoxetine treatment for each genotype (see Figure in https://phenome.jax.org/measureset/38005) for the DS phenotype comparing treatment effects between the different strains. In such a case, a linear contrast could have been performed for statistically based inference as follows:

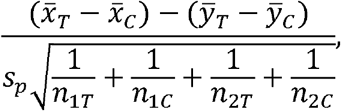

Where 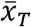,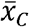 denote the mean size at the treatment and control groups of the first genotype, respectively, and *n*_1*T*_, *n*_1*C*_ denote their sample sizes. For the second genotype, *ȳ_T_*, *ȳ_C_* denote the mean size at the treatment and control groups, respectively and *n_2T_*, *n_2C_* denote their sample sizes.

The second order interaction of treatment by genotype is the fixed parameter we try to estimate. All other second order interactions, namely treatment-by-lab and the genotype-by-lab cancel out in the contrast. What remains is the third order interaction treatment-by-genotype-by-lab (TxGxL) that specifies the random contribution of the laboratory on the measurement for a specific genotype when treated and another one when not. Since there are 4 of these, and they are independent, they increase the variance by 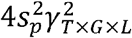,

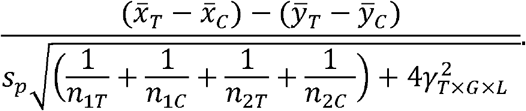

The degrees of freedom are recalculated again according to the Satterthwaite formula. As previously mentioned, we do not have an outer (independent) source to estimate the size of this interaction. Therefore, we estimate it using the data collected by our 3-lab experiment, and we apply it on the t-tests of the original Wiltshire2 study. Due to the lack of a third-party estimate, we are unable to demonstrate the power and type-I error of the tool, but merely assess the robustness of the original study results to the proposed adjustment.

#### 4.5.3 Displaying the results

To study the effect of the adjustment we treat our 3-lab replication results as our “ground truth”, where a statistically significant difference is considered a “replicated difference” (Table 1 first column). For each difference between two strains in each phenotype we then examine if (second column) this difference was significant in the original single-lab study in the MPD, and if (third column) it is still significant after correcting it using the GxL-adjustment calculated in the IMPC (Kafkafi et al., 2017). The six possible combinations are denoted by categories A–F (fourth column), with their interpretations (fifth column).

We derive the proportion of the non-replicable differences (categories D+E+F) that were inappropriately discovered by the original analysis study (D+E). We also derive the same proportion after both readjusted (D). As to loss of replicable discoveries the proportion of replicated ones discovered by the original analysis and after its adjustment are given by the last two lines in the table. (Note that if we treat replicated difference as ‘true’ and non-replicated difference as false we discuss Type-I error and power with and without adjustment).

#### 4.5.4 Computational details and data availability

Statistical analysis was done in R version 4.1.1 (2021-08-10) (R Core Team, 2021). We use the packages “lme4” (Bates et al. 2015), “nlme” (Pinheiro et al. 2021) and “multcomp” (Hothorn et al. 2008) to perform multi-lab analysis via Restricted Maximum Likelihood (REML) and pairwise comparisons. Figures shown in this paper are produced with the package “ggplot2” (Wickham, 2016).

Raw data files and R code for reproducible research is available online at GitHub. URL: https://github.com/IJaljuli/Improving-replicability-using-interaction-2021.

## Acknowledgements

This work was supported by the US–Israel Binational Science Foundation—US National Science Foundation (BSF-NSF 2016746) (NIH-NSF DA045401). The authors are grateful to Dr. Lior Bikoveski at the Myers Neuro-Behavioral Core Facility for his continuous support with our mouse behavioral studies at the Faculty of Medicine, Tel Aviv University. We gratefully acknowledge support from *P30 CA034196* and the contributions of Zoe Bichler, Laura Anderson, Torrian Green, Meaghan Dyer, Veronica Knickerbocker, and the Center for Biometric Analysis Service for expert assistance with phenotyping conducted at The Jackson Laboratory. We would also like to thank the JAX Computational Sciences team that provided software development, quality assurance, and software engineering for the GxL Replicability Adjuster tool in MPD, including Georgi Kolishovski, Hao He, Vivek Philip, Dave Walton, Stephen Grubb, Jake Emerson, Matt Dunn, Vinita Sinha, and Beena Kadakkuzha.

## Supplementary Material

**Table S1:** Timelines of animals and tests participating in the IMPC database (EUMODIC Pipeline 2), in the 5 replicated studies in the MPD database, and in the three labs participating in the experiment: TAUL, TAUM and JAX. Tests in Bold letters are common to all databases and labs.

| Week after birth | IMPC (EUMODIC Pipeline 2) [ESLIM_002] | MPD: Crabbe4 | MPD: Tarantino2 | MPD: Tordoff3 | MPD: Wiltshire2 | MPD: Crowley1 | TAUL | TAUM | JAX |
| --- | --- | --- | --- | --- | --- | --- | --- | --- | --- |
| 5 |  | Shipping | Shipping |  | Shipping |  | Moving |  | Shipping |
| 6 |  | Shipping | Shipping |  | Begin fluoxetine |  | Begin fluoxetine | Begin fluoxetine | Begin fluoxetine |
| 7 |  | **GS** | Sham surgery |  |  |  |  |  |  |
| 8 |  | **GS** |  | **BW** |  | Shipping | **GS; BW** | **GS; BW** | **GS; BW** |
| 9 | **OF** large arena<br>**GS; BW;**<br>Modified SHIRPA | **GS** |  | **BW;**<br>preference for CaCl<sub>2</sub> | **TS** | **BW** | **GS; BW**<br>**OF** females: large arena<br>**OF** males: small arena | **GS; BW**<br>**OF** females: large arena<br>**OF** males: small arena | **GS; BW;**<br>**OF** females: large arena |
| 10 | Rotarod | **GS** | **OF** females: Large arena | **BW** | **TS; OF** males, small arena |  | **OF** females: large arena<br>**OF** males: small arena | **OF** females: large arena<br>**OF** males: small arena | **OF** females: large arena |
| 11 | Acoustic startle & PPI | **GS** | **OF** females: Large arena |  | **OF** males, small arena |  | **OF** females: large arena<br>**OF** males: small arena<br>**TS** | **OF** females: large arena<br>**OF** males: small arena<br>**TS** | **OF** males: large arena; **TS** |
| 12 | Hot Plate | **GS** | **OF** females: Large arena |  |  |  | **TS;**<br>**OF** males, large arena | **TS;**<br>**OF** males, large arena | **OF** males: large arena; **TS** |
| 13 | Blood tests |  |  |  |  |  | **OF** males, large arena | **OF** males, large arena | **OF** males: large arena |

**Table S2:** OF parameters used in different mouse groups in each of the three labs, in the IMPC database and in the MPD studies.

| Lab | Gender | Age | Treatment | Arena Size | Tracking method (system, company) | Periphery definition | Session duration |
| --- | --- | --- | --- | --- | --- | --- | --- |
| TAUM | Females | 10 weeks | Control | 42 cm | Video (Ethovision, Noldus) | 10 cm | 10, 20 min |
| TAUL | Females | 9-11 weeks | Control | 42 cm | Video (Ethovision, Noldus) | 10 cm | 10, 20 min |
| JAX | Females | 9-11 weeks | Control | 40 cm | Photobeam (Omnitech) | 8 cm | 10, 20 min |
| IMPC | Females | 8 weeks | Control | 40-44 cm | Video (Ethovision, Noldus) | 16 cm | 20 min |
| Tarantino2 | Females | 11 weeks | Control | 42 cm | Photocell (Versamax, AccuScan, Columbus) | 9.57 cm | 10 min |
| TAUM | males | 10 weeks | Control | 27 cm | Video (Ethovision, Noldus) | 8 cm | 10, 20 min |
| TAUL | males | 9-11 weeks | Control | 27 cm | Video (Ethovision, Noldus) | 8 cm | 10, 20 min |
| JAX | males | 9-11 weeks | Control | 40 cm | Photobeam (Omnitech) | 8 cm | 10, 20 min |
| IMPC | males | 8 weeks | Control | 40-44 cm | Video (Ethovision, Noldus) | 16 cm | 20 min |
| Wiltshire2 | males | 10-11 weeks | Control | 27.3 cm | Photobeam (MED-OFA-MS, Med Associates) | 6.8 cm | 10 min |
| TAUM | males | 9-11 weeks | Fluoxetine | 27 cm | Video (Ethovision, Noldus) | 8 cm | 10, 20 min |
| TAUL | males | 9-11 weeks | Fluoxetine | 27 cm | Video (Ethovision, Noldus) | 8 cm | 10, 20 min |
| JAX | males | 9-11 weeks | Fluoxetine | 40 cm | Photobeam (Omnitech) | 8 cm | 10, 20 min |
| TAUM | males | 11-12 weeks | Control | 42 cm | Video (Noldus) | 10 cm | 10, 20 min |
| TAUL | males | 11-12 weeks | Control | 42 cm | Video (Noldus) | 10 cm | 10, 20 min |
| TAUM | males | 11-12 weeks | Fluoxetine | 42 cm | Video (Noldus) | 10 cm | 10, 20 min |

**Table S3:** Estimated GxL factors γ for different subgroups and endpoints including those estimated from 3-way analysis of drug treatment genotype and labs and those estimated from IMPC experiments.

| Test | | Phenotype | Session length | Arena size | 3-lab GxL factor ($\gamma$) | IMPC GxL factor | 3-lab GxL factor for Fluoxetine |
| --- | --- | --- | --- | --- | --- | --- | --- |
| Open Field | Female | Dist. Traveled | 10min | Small |  |  |  |
|  |  |  |  | Large | 0.898 | 0.360 |  |
|  |  |  | 20min | Small |  |  |  |
|  |  |  |  | Large | 0.926 |  |  |
|  |  | % Cent. Time | 10min | Small |  |  |  |
|  |  |  |  | Large | 0.595 | 0.459 |  |
|  |  |  | 20min | Small |  |  |  |
|  |  |  |  | Large | 0.501 |  |  |
|  | Male | Dist. Traveled | 10min | Small | 0.974 | 0.325 | 0.206 |
|  |  |  |  | Large | 1.250 |  | 0.425 |
|  |  |  | 20min | Small | 1.254 |  | 0.346 |
|  |  |  |  | Large | 1.325 |  | 0.380 |
|  |  | % Cent. Time | 10min | Small | 0.00 | 0.133 | 0.490 |
|  |  |  |  | Large | 0.480 |  | 0.527 |
|  |  |  | 20min | Small | 0.325 |  | 0.423 |
|  |  |  |  | Large | 0.425 |  | 0.365 |
| Grip Strength | Female |  |  |  | 0.537 | 0.719 |  |
|  | Male |  |  |  | 0.541 | 0.543 | 0.688 |
| Body Weight | Female |  |  |  | 0.545 | 0.266 |  |
|  | Male |  |  |  | 0.611 | 0.21 | 0.610 |
| Tail Suspension | Female |  |  |  | 0.400 |  |  |
|  | Male |  |  |  | 0.152 |  | 0.591 |

**Table S4:** Results of naïve and GxL-adjusted genotypic differences for all phenotypes in the MPD experiments, using GxL factors estimated from the IMPC standardized data.

| category | interpretation | No.<br>cases | %<br>cases |
| --- | --- | --- | --- |
| A | Replicable discoveries, approved by adjustment | 20 |  |
| B | Replicable discoveries, missed by adjustment | 10 |  |
| C | Replicable discoveries missed by both | 3 |  |
| D | Non-replicable, insufficient adjustment | 6 |  |
| E | Non-replicable, prevented by adjustment | 24 |  |
| F | Non-replicable by both | 29 |  |
| $(D+E)/(D+E+F)$ | Proportion of non-replicable discoveries | 30/59 | 50.9% |
| $(D)/(D+E+F)$ | Proportion of non-replicable discoveries, GxL-adjusted | 6/59 | 10.2% |
| $(A+B)/(A+B+C)$ | Power to discover replicable discoveries | 30/33 | 90.9% |
| $(A)/(A+B+C)$ | Power to discover replicable discoveries, GxL-adjusted | 20/33 | 60.6% |

**Figure S1:**
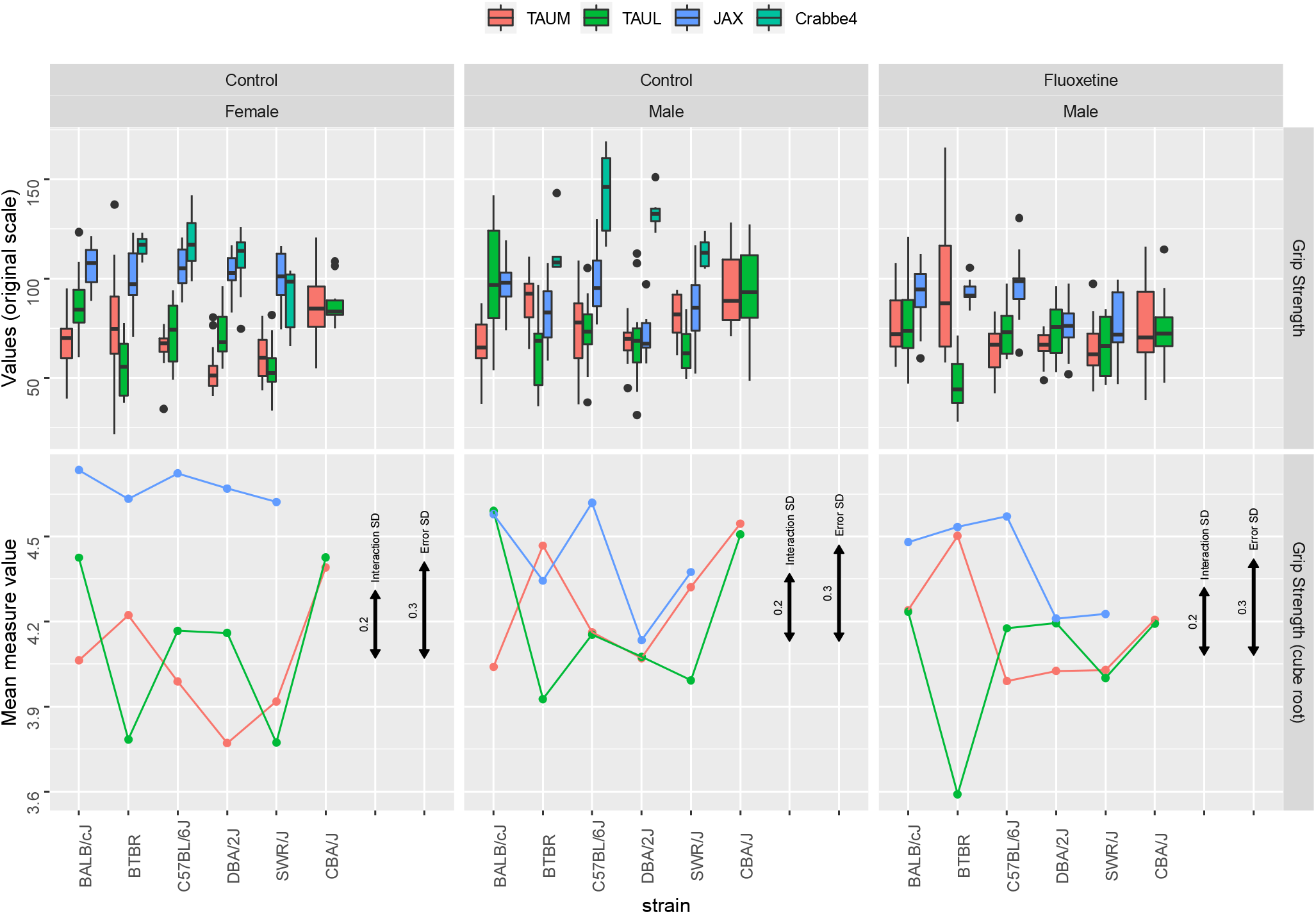
Forepaw peak strength results in the Grip Strength (GS) test, in the 3-lab experiment and the MPD study Crabbe4, using boxplots (top) and genotype means after raising to the power of 1/3 transformation (bottom), in females (left), males (center) and fluoxetine-treated males (right). Black error bars represent the interaction SD and the within-group error SD.

**Figure S2:**
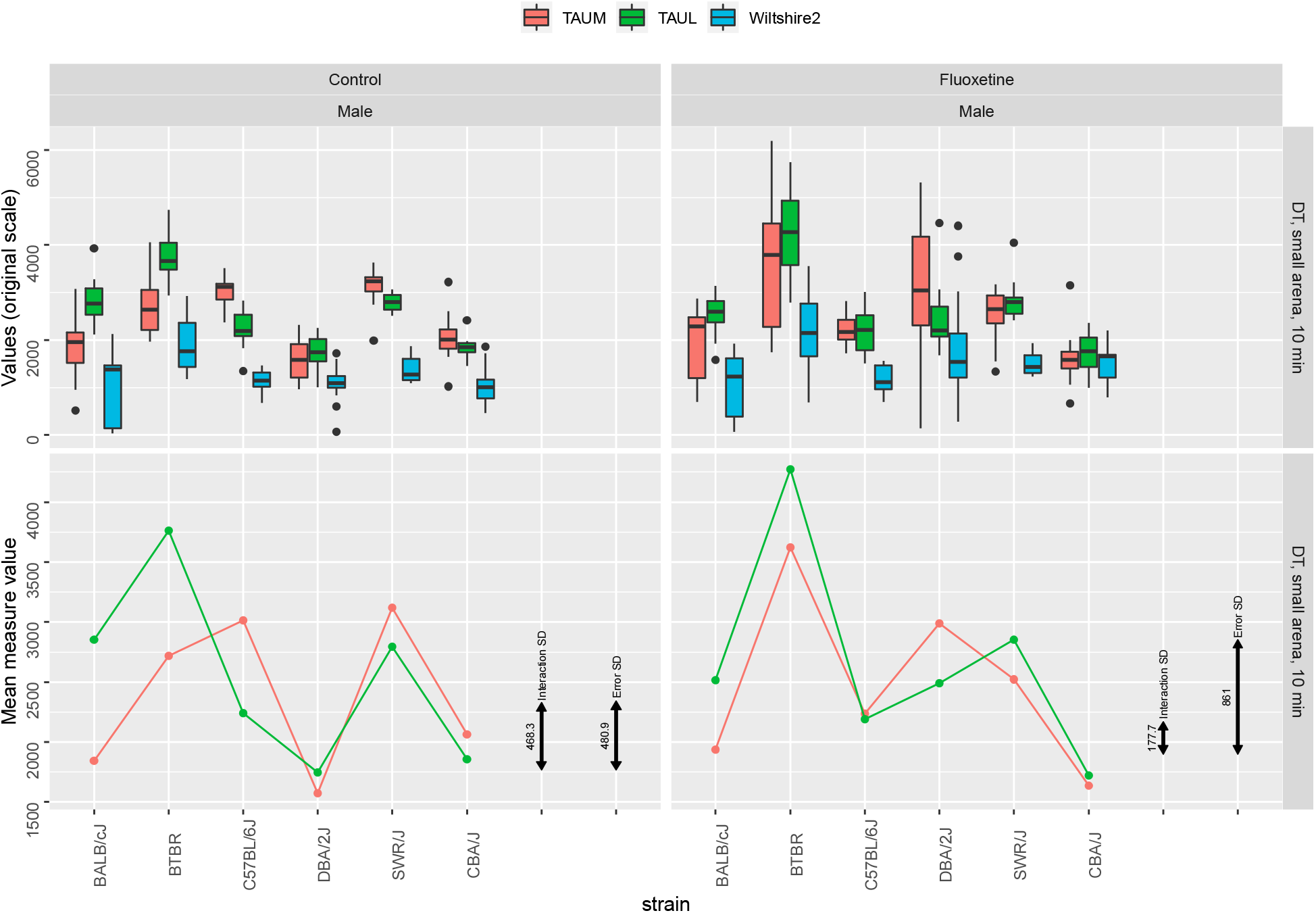
Distance Traveled (DT) in the Open Field (OF) test, in a small arena in 10 minutes, in the 3-lab experiment and the MPD study Crabbe4, using boxplots (top) and genotype means (bottom), in females (left), males (center) and fluoxetine-treated males (right). Black error bars represent the interaction SD and the within-group error SD.

**Figure S3:**
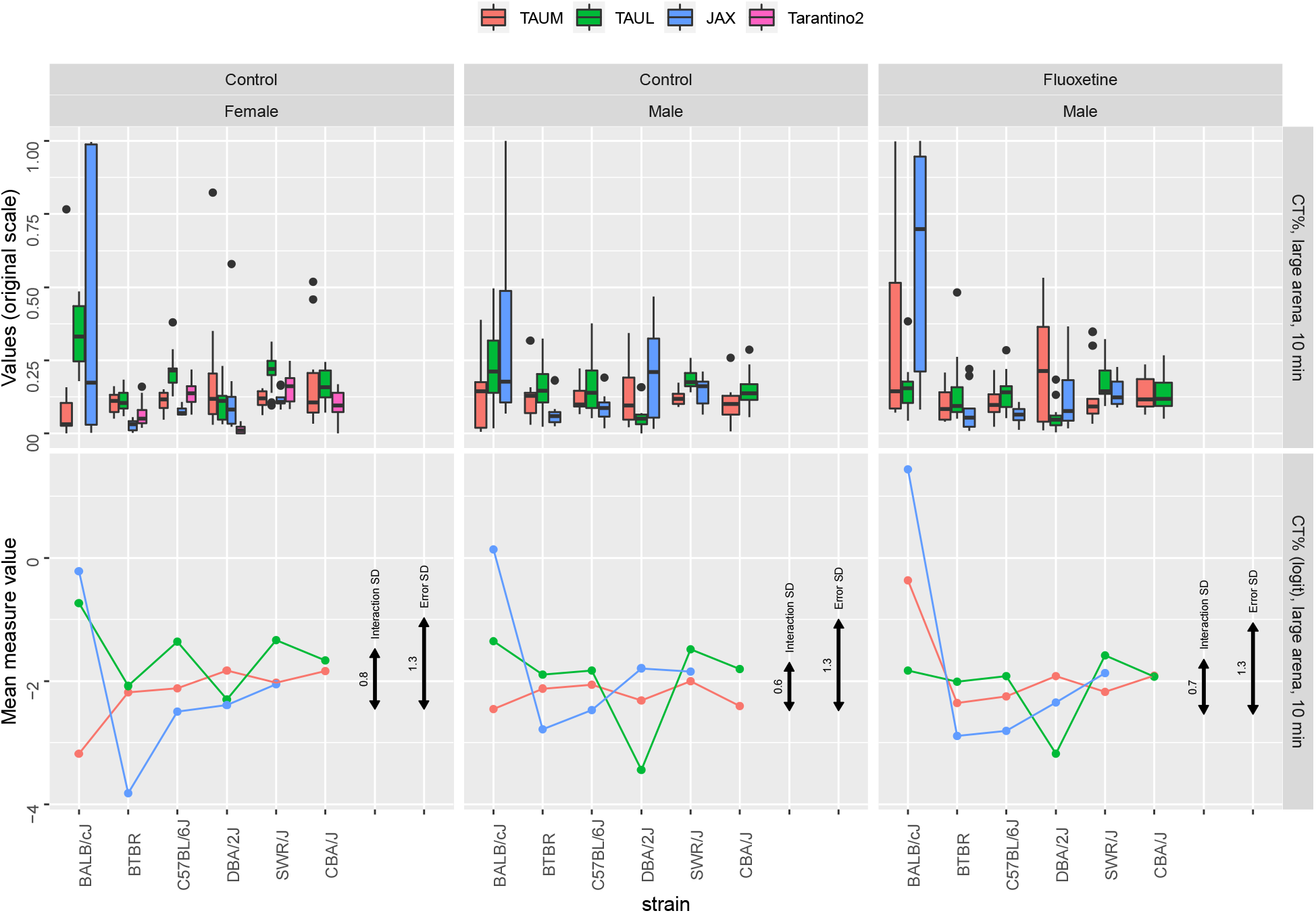
% Center Time % in the Open Field (OF) test, in a large arena for 10 minutes, in the 3-lab experiment and the MPD study Tarantino2, using boxplots (top) and genotype means after logit transformation (bottom) in the three laboratories, in females (left), males (center) and fluoxetine-treated males (right). Black error bars represent the interaction SD and the within-group error SD.

**Figure S4:**
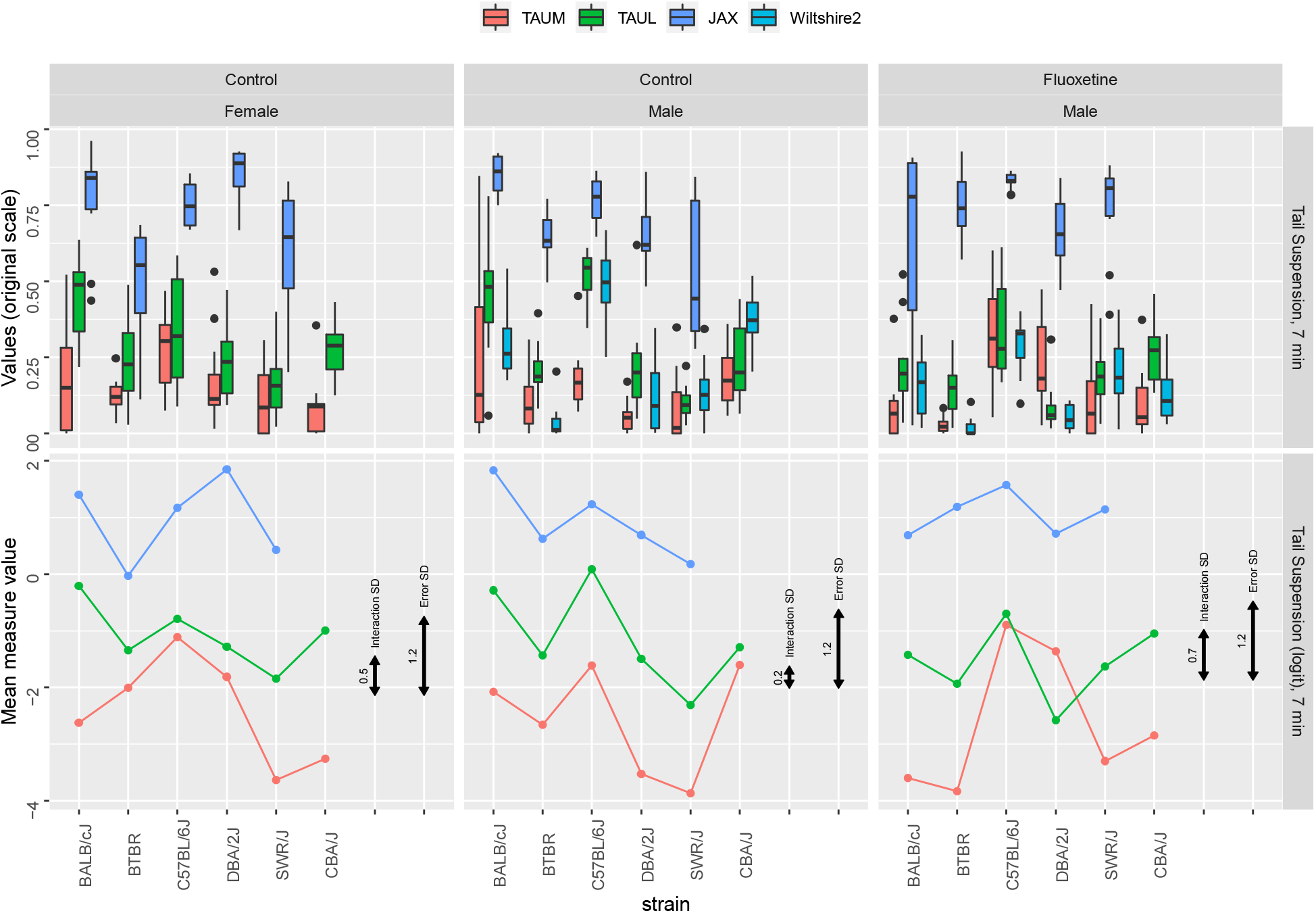
% time spent in immobility during 7 min in the Tail Suspension (TS) test, in the 3-lab experiment and the MPD study Wiltshire2, using boxplots (top) and genotype means after logit transformation (bottom), in females (left), males (center) and fluoxetine-treated males (right). Black error bars represent the interaction SD and the within-group error SD.

**Figure S5:**
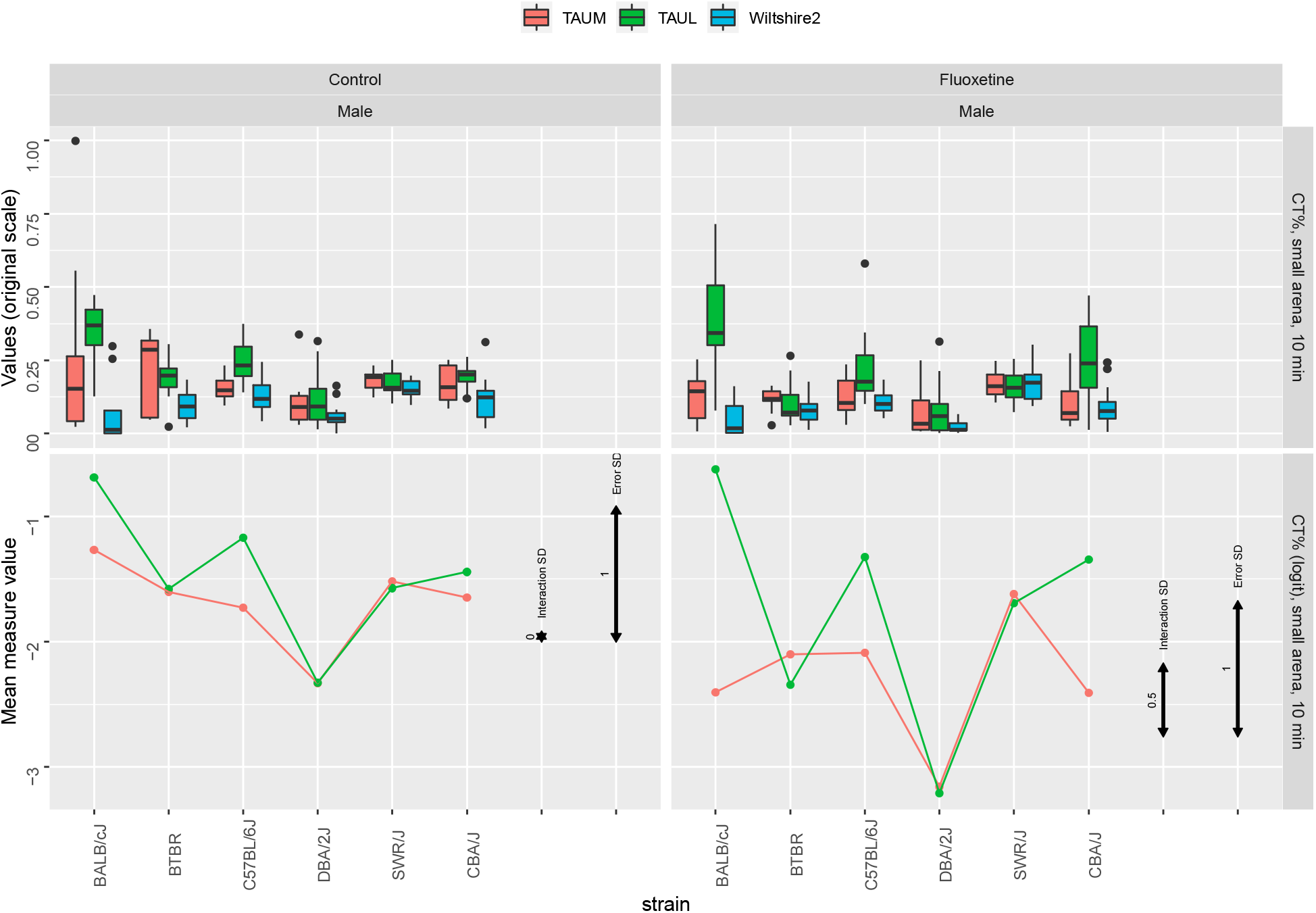
% Center Time % (CT) results in the Open Field (OF) test, in a large arena for 10 minutes, in the 3-lab experiment and in the MPD study Wiltshire2, using boxplots (top) and genotype means after logit transformation (bottom), in females (left), males (center) and fluoxetine-treated males (right). Black error bars represent the interaction SD and the within-group error SD.

## Notes

https://github.com/IJaljuli/Improving-replicability-using-interaction-2021

### Competing Interest Statement

The authors have declared no competing interest.

### Summary of Updates

Mainly language editing

